# Deviation from the Residue-Based Linear Free Energy Relationship Reveals a Non-Native Structure in Protein Folding

**DOI:** 10.1101/2022.01.07.475459

**Authors:** Daisuke Fujinami, Seiichiro Hayashi, Daisuke Kohda

## Abstract

Multiprobe measurements, such as NMR and hydrogen exchange study, can provide the equilibrium constant K and kinetic rate constant k of the structural changes of a polypeptide on a per-residue basis. We previously found a linear relationship between residue-specific log K values and residue-specific log k values for the two-state topological isomerization of a 27-residue peptide. To test the general applicability of the residue-based linear free energy relationship (rbLEFR), we performed a literature search to collect residue-specific equilibrium and kinetic constants in various exchange processes, including protein folding, coupled folding and binding of intrinsically disordered peptides, and structural fluctuations of folded proteins. The good linearity in a substantial number of log-log plots proved that the rbLFER holds for the structural changes in a wide variety of protein-related phenomena. Protein molecules quickly fold into their native structures and change their conformations smoothly. Theoretical studies and molecular simulations advocate that the physicochemical basis is the consistency principle and the minimal frustration principle: Non-native structures/interactions are absent or minimized along the folding pathway. The linearity of the residue-based free energy relationship demonstrates experimentally the absence of non-native structures in transition states. In this context, the hydrogen exchange study of apomyoglobin folding intermediates is particularly interesting. We found that the residues that deviated from the linear relationship corresponded to the non-native structure, which had been identified by other experiments. The rbLFER provides a unique and practical method to probe the dynamic aspects of the transition states of protein molecules.

**Highlights:**

- A collection of equilibrium and kinetic constants of structural changes of proteins
- Residue-based linear free energy relationship widely holds between the two constants
- rbLFER indicates the absence of non-ground state structures in transition states
- rbLFER is an experiment proof of the consistency principle of protein folding
- Deviations from the linear relation suggest special structures in transition states

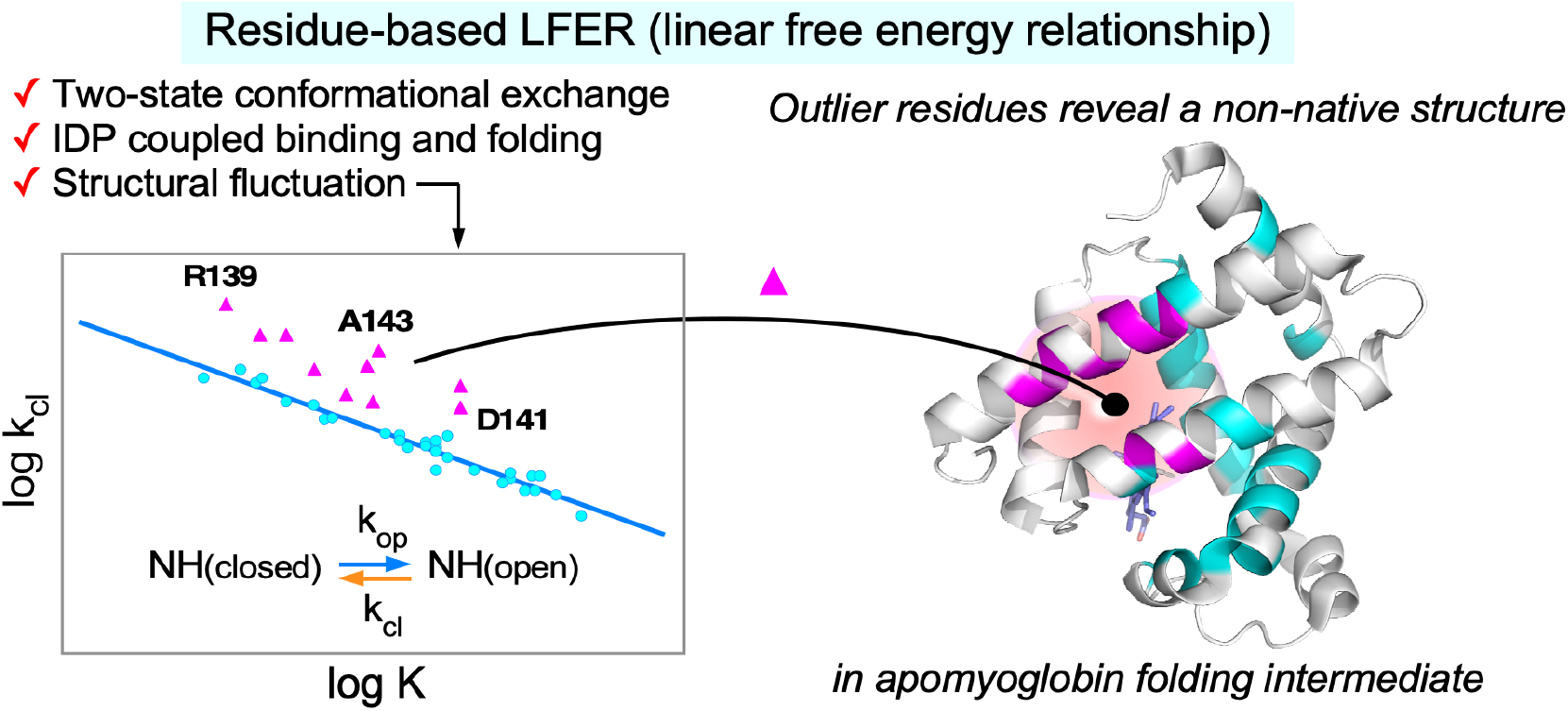

## Introduction

Protein molecules quickly fold into their thermodynamically stable and biologically active structures. Protein folding is a property acquired by evolution and is highly optimized for each amino acid sequence. Nobuhiro Gō proposed the ‘consistency principle’ theory in 1983[1]. The consistency principle of protein folding is the physicochemical basis for a perfectly smooth surface of the funnel-like energy landscape to materialize the smooth folding of a polypeptide chain. It advocates the absence of non-native structures or interactions in every moment on the folding paths from any random initial structure. In 1987, Bryngelson and Wolynes proposed a more realistic version of the consistency principle, based on the theory of spin glasses[2]. Through evolutionary selections, the surface of the funnel-like landscape has been sculpted to become smooth by minimizing the frustration of tertiary interactions, but remains rugged to the minimum necessary for satisfying the functionality requirements, such as enzymatic activities and allostery of proteins[3,4]. The consistency principle and the minimum frustration principle have been widely supported by many molecular simulations. In 2013, the refolding trajectories of small proteins obtained by all-atom MD simulations with explicit water solvent were analyzed from the perspective of folding mechanisms[5]. In the successful refolding paths, native contacts were formed at higher probabilities than non-native contacts, strongly supporting the viewpoint of the consistency principle.

NMR provides local information around nuclei at an atomic resolution. We used NMR to determine the residue-specific equilibrium constants and residue-specific kinetic rate constants of a polypeptide chain in a two-state dynamic exchange, and discovered a linear relationship in the double logarithmic plot of the two parameters[6]. This type of plot is called a REFER (rate equilibrium free energy relationship) plot, and the linear relationship in this plot is generally referred to as the Linear Free Energy Relationship (LFER). LFER is widely observed in many chemical and biological fields and used to investigate the transition states of reaction and exchange phenomena[7–12]. Importantly, all known types of LFER are composed of data points obtained from single probe measurements. Data points, (K, k), are derived from different molecules by independent measurements, in which a perturbation is a difference in the chemical structure (including amino acid sequence), or from one molecule by repeated measurements under different perturbed conditions, such as variations in the solvent. The interpretation of the results always requires the assumption that the perturbations do not affect the transition paths. In contrast, the LFER we found is derived from many amino acid residues of one polypeptide chain under a single set of conditions. The “perturbation” is the different positions of residues as the NMR probe. In this respect, we refer to this new type of LFER as residue-based LFER (rbLFER). Obviously, rbLFER is not affected by unintended changes of the transition paths.

Residue-based LFER was discovered in the NMR study of a 27-residue bioactive peptide, nukacin ISK-1[6]. The two sets of ^1^H−^15^N amide cross peaks in the ^1^H−^15^N HSQC (heteronuclear single quantum coherence) spectrum indicated the existence of two states in solution, termed states A and B[13]. The interconversion rate was slow on the second time scale, reflecting the high energy barrier, 16 k_B_T. The peak intensity ratio of the two cross peaks of each residue in the HSQC spectrum provides an equilibrium constant, K. Unexpectedly, we found that the K values varied from one residue to another[14]. Therefore, we determined the exchange rates, k (forward) and k’ (backward), using EXSY (exchange spectroscopy)[6]. We found that both the residue-specific k and k’ values also varied. Subsequently, we examined the relationship between the residue-specific thermodynamic and kinetic terms and found a linear relationship between log k and log K in the REFER plot. In principle, the equilibrium constant and the kinetic rate constants are independent, but the residue-based LFER suggested a hidden relationship between the transition state and the two ground states over the entire polypeptide chain. Our present interpretation is that the physicochemical basis of the smooth transition between the two states emerges as the linear relationship in the REFER plot of nukacin ISK-1.

Here, we performed a literature search to collect residue-specific equilibrium and kinetic constants of proteins and generated REFER plots. A substantial number of the REFER plots exhibited good linearity, which is, in our opinion, a direct result of the consistency principle of protein conformational changes. Among the successful cases, the HX (hydrogen exchange) study of apomyoglobin folding intermediates is particularly interesting[15]. The majority of the amino acid residues are on a straight line in the REFER plot, but some residues deviate from the line. We found that these outlier residues precisely correspond to the non-native structures in the apomyoglobin folding intermediates, which were first discovered a decade- and-a-half ago[16,17]. The excellent agreement between the two independent analyses demonstrates that rbLFER can provide the structural aspects of the dynamic properties of protein molecules.

## Results

### Residue-based LFER examples

Extensive literature searches led to the collection of reports on residue-specific equilibrium constants and residue-specific kinetic constants of proteins (Table 1). We recovered the data from the tables and figures and generated the corresponding REFER plots (Fig. 1). The techniques used and the phenomena under consideration are quite diverse. The first category (including nukacin ISK-1) is the EXSY NMR analyses of slow exchanges between two sets of cross peaks (Fig. 1A and 1B)[6,18,19], and the second one is the relaxation dispersion (RD) NMR analyses of the binding of an IDP (intrinsically denatured polypeptide) to a target protein (Fig. 1C)[20,21]. In the case of an IDP, only one set of cross peaks was observed due to the fast exchange rates, but the information on the exchange process can be extracted from the averaged cross peaks. The same situation is applied to the structural fluctuations of native-state proteins studied by RD NMR (Fig. 1D)[22–27]. The HX experiment also provides information on the structural fluctuations of native proteins (Fig. 1E)[28–31]. Note that the NMR in the HX studies is used for the quantitation of the proton occupancy and does not directly observe the HX phenomena. Indeed, mass spectrometry is applicable in place of NMR. Overall, the majority of the REFER plots showed residuebased linear relationships.

**Table 1.**
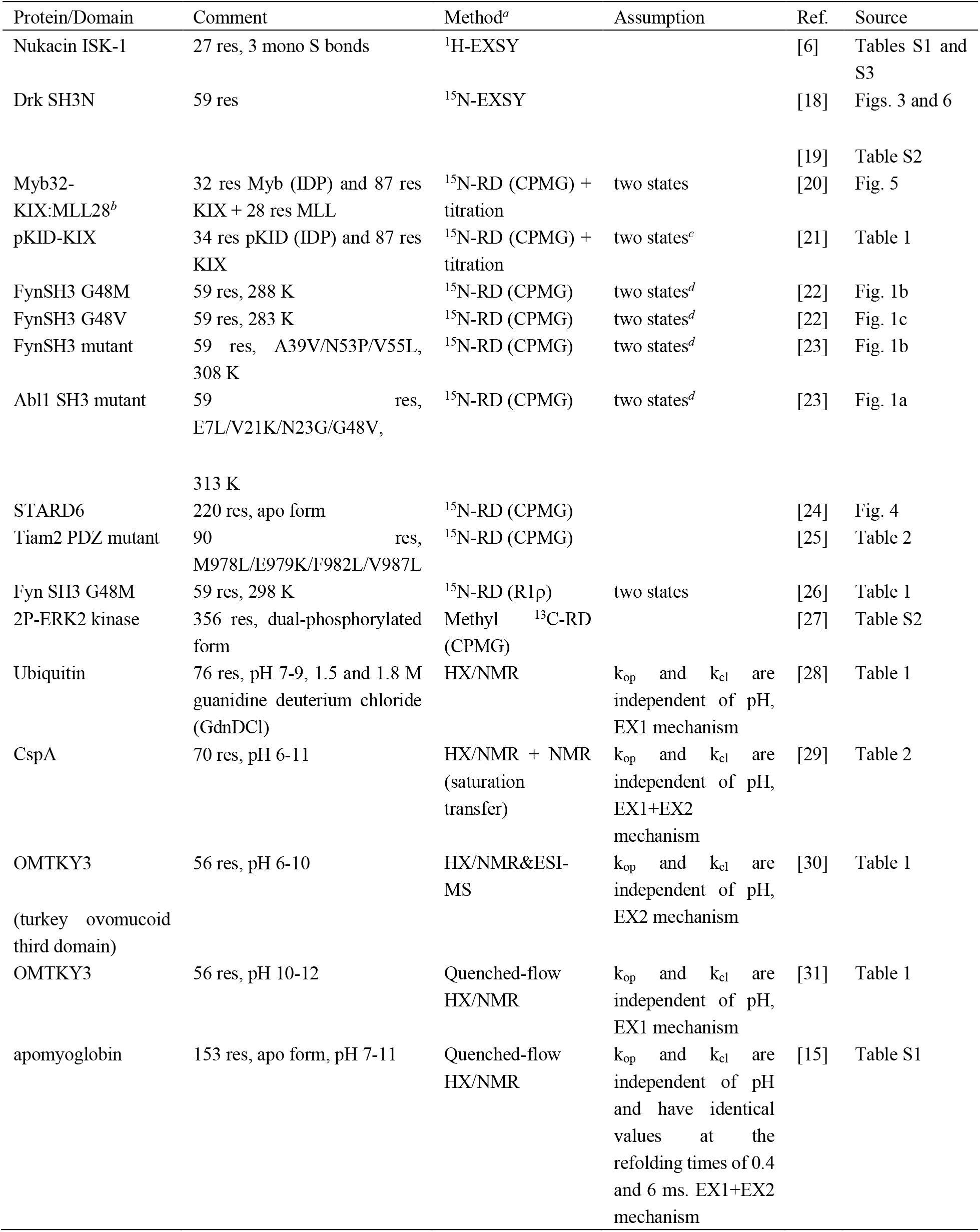
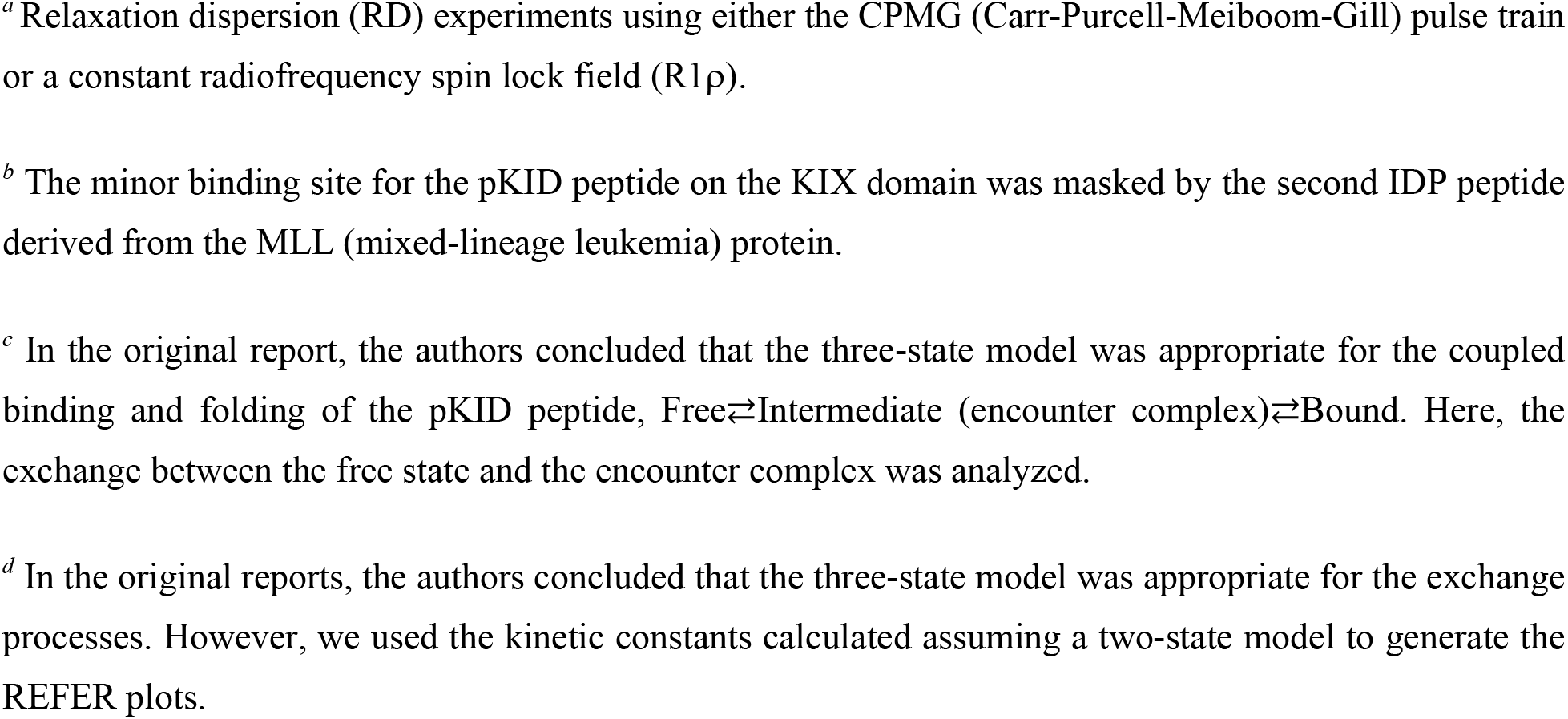
Proteins used for the generation of REFER plots

**Fig. 1.**
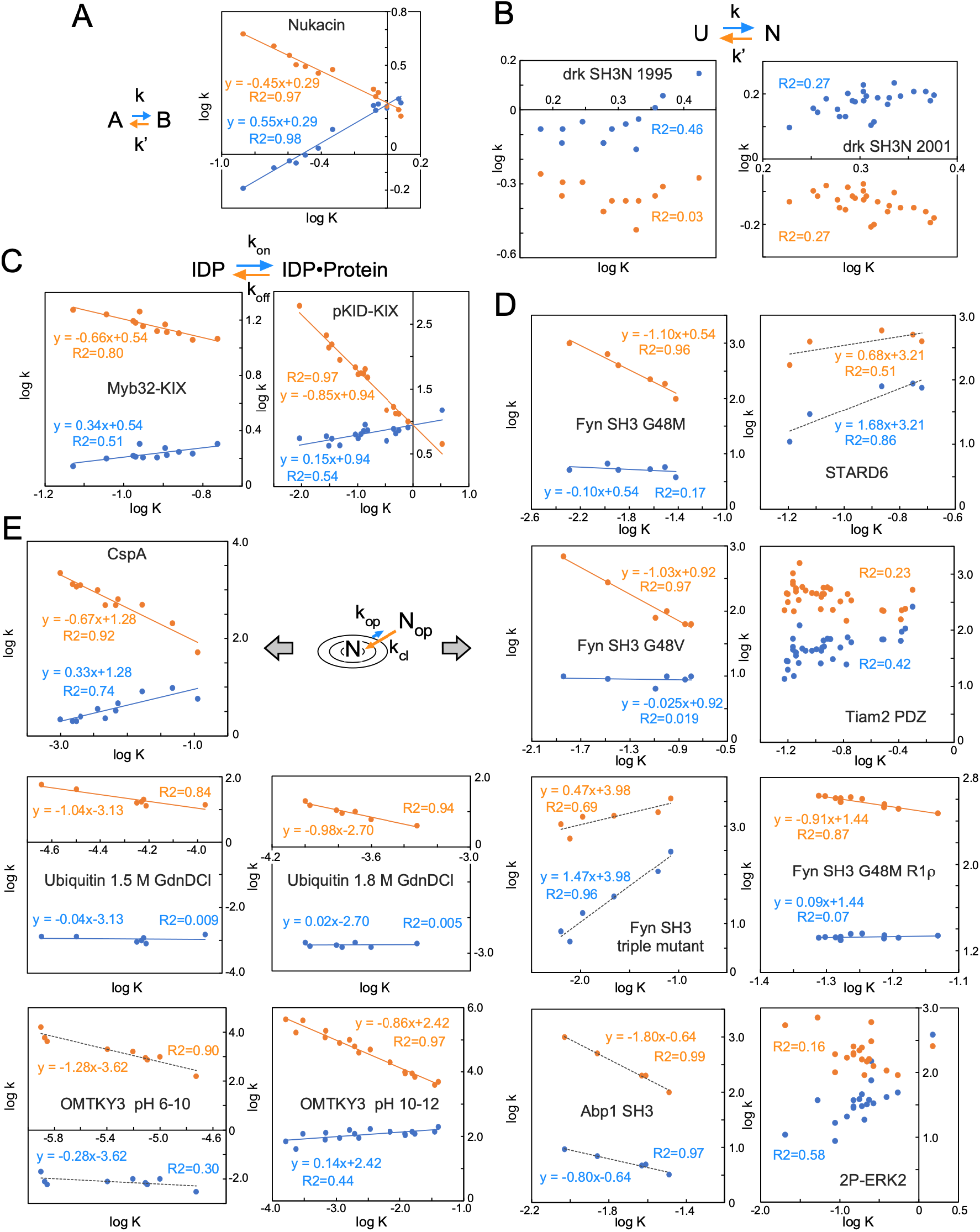
Residue-based REFER plots based on the literature data listed in Table 1. (A) Nukacin ISK-I. (B) N-terminal SH3 domain from Drosophila drk protein (drk SH3N). The drk SH3N is in an exchange between unstructured state U and native folded state N. There are different data sets from two reports. (C) Myb32 and pKID are intrinsically disordered polypeptides that bind to the KIX domain. (D) Structural fluctuations of native states were investigated by the relaxation dispersion (RD) NMR method. The SH3 domains are derived from the Fyn and Abp1 proteins. The mutations in the SH3 domains markedly increased the folding rate despite their destabilization of the folding state, and have suitable properties for the RD studies[36]. The STARD (steroidogenic acute regulatory-related lipid transfer domain) from the STARD6 protein. The PDZ (PSD-95/Dlg/ZO-1) domain from the Tiam2 (T cell lymphoma invasion and metastasis 2) protein. A dual-phosphorylated (2P-ERK2) form of the ERK2 (extracellular signal-regulated kinase 2) protein. (E) Structural fluctuations of native states investigated by the hydrogen exchange method. There are two measurement conditions for ubiquitin and OMTKY3 (turkey ovomucoid third domain 3). In (A)-(E), the least-square lines and data points associated with the forward direction are colored blue, and those associated with the backward direction are orange. The least-square lines with interpretable slopes between 0 and 1 are depicted as solid, colored lines, whereas those with uninterpretable slopes less than 0 or greater than 1 are depicted as dashed black lines. No least-square lines are drawn if the correlations are considered insignificant. The pairs of arrows show the interconversion processes between two states and the structural fluctuations of native-state proteins. The concentric circle represents a basin-shaped energy landscape of the structural fluctuations around the native state, N, in equilibrium with the open state, N_op_.

Here, we discuss the interpretation of the slopes of the REFER plots, by focusing on the forward direction (right-pointing arrows) depicted by the blue lines and their associated data points (Fig. 1). The slope represents the structural and energetical similarity between the starting ground state and the transition state, and hence the value must be between 0 and 1. In the case of structural fluctuations of the native states (Fig. 1D and 1E), the blue lines are nearly horizontal, and the slopes are almost zero in many instances. This situation indicates the high similarity of the transition state to the native state. This is a convincing result, considering that the interactions that stabilize the native state must be disrupted at the beginning of the fluctuation. The least-square lines with negative slopes and slopes greater than 1 are difficult to interpret. These cases are depicted by dashed lines. In some cases, particularly for large proteins, the data points are scattered, and no least-square lines are shown. Understandably, the REFER plots based on the literature data, including those with interpretable slopes, must be properly assessed in the future.

### Alternative representation of residue-based LFER

The physical interpretation of the REFER plot is clear: the two axes, log K and log k, are proportional to the changes in free energy terms. However, the two axes are not independent due to the equation, log K = log k − log k’, which tends to exaggerate the linearity of the REFER plot (see the next section). Therefore, the log k vs. log k’ plot is suitable for the assessment of rbLFER. Figure 2 shows the type classifications of log k vs. log k’ plots, and Figure 3 shows the log k vs. log k’ plots of the real examples listed in Table 1. The classification types are defined according to the distribution pattern of the data points. Type N is referred to as the negative correlation between log k and log k’, which leads to the blue least-square lines with the slope of 0 < ρ < 1 in the REFER plot (cf. equation 9 in the previous paper[6]). In extreme cases, when the distribution of data points in the log k vs. log k’ plot has a flattened shape, the least square lines have the slope of 0 or 1. According to the orientation of the oval-shaped distribution, vertical and horizontal, their types are defined as V and H, respectively. Since either log k or log k’ is rather constant, the two log terms are uncorrelated in types V and H. Consequently, the Pearson’s correlation coefficient R is zero in the log k vs. log k’ plot, and one of the two least-square lines with a zero slope has an almost zero R^2^-value in the REFER plot. This fact indicates that the R and R^2^ values in the two plots are not always good indicators of rbLFER. Instead, 95% confidence ellipses are drawn to quantify the flat distributions (Fig. 3A-E). The closer flatness of the distribution to 1 reflects the higher linearity of the rbLFER. Next, we consider the case of positive correlations between log k and log k’. If the degree of variation of log k’ is larger than that of log k, the type is P, and if the degree of variation of log k’ is smaller than that of log k, the type is P’. Because the slopes of the blue least-square lines become negative or greater than 1 according to the type, the slopes are not physically meaningful. Such anomalous slope values occur if the exchange process cannot be described by a simple two-state model. For example, the ^15^N relaxation dispersion studies of various mutants of the Fyn SH3 domain revealed the presence of about 1% of the transient intermediate state I in the three-state model, N⇄I⇄U[22,23,26]. Even though the percentage is small, the assumption of the two-state model is not strictly valid, and thus the log k vs. log k’ plots do not always show meaningful rbLFER properties (Fig. 1D). Another possible cause is unintended measurement biases. The final type is nr (no relation). This is probably due to large measurement errors or accidental problems. For example, in the case of drk SH3N (Fig. 1B), the NMR spectra were recorded at pH 6. Under near-neutral conditions, the large exchange rate of amide protons with bulk water might reduce the measurement accuracy, despite the use of special NMR pulse sequences[18]. In the nukacin ISK-1 experiments, we used a low pH (pH 3.5) to slow down the water exchange and the P analysis to identify problematic data points affected by the exchange with water[6].

**Fig. 2.**
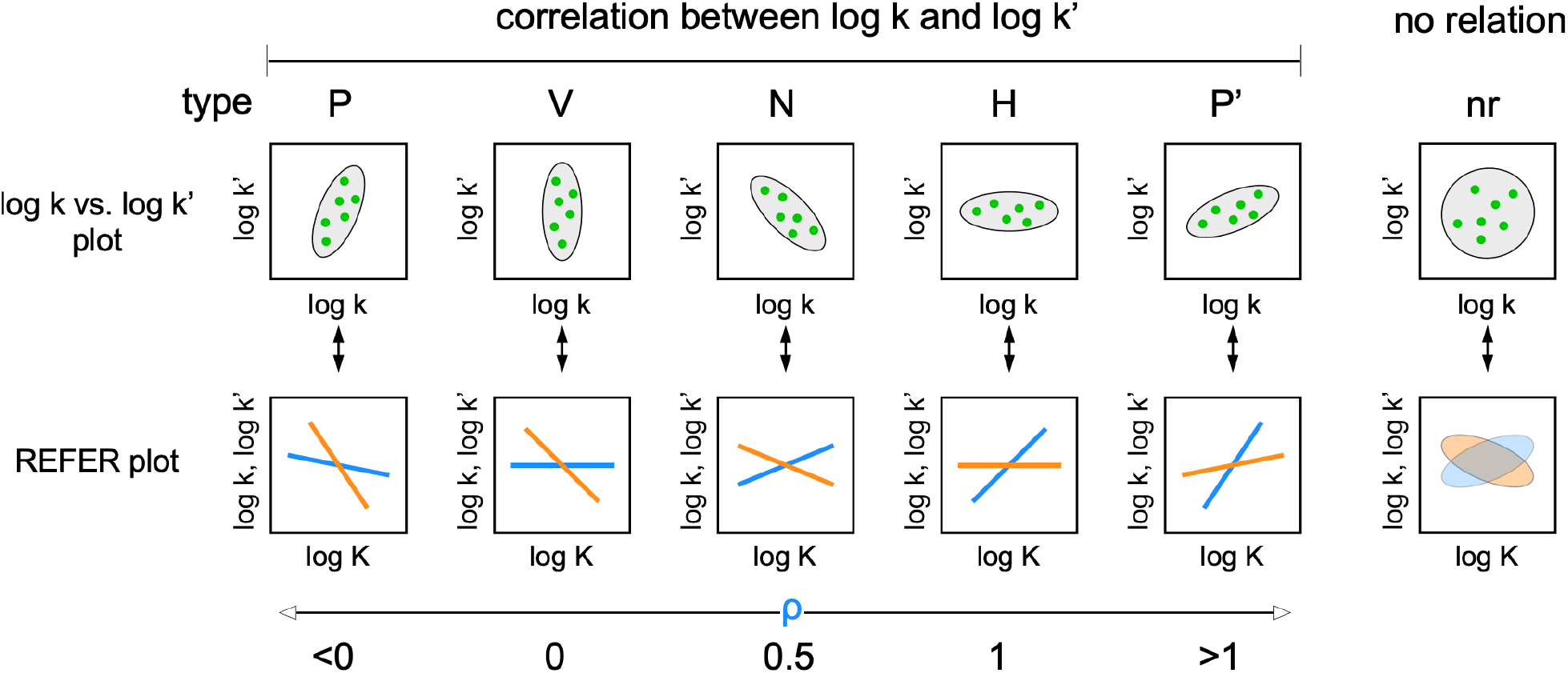
Relationships between log k and log k’ as the basis of rbLFER. Several types are defined according to the data point distribution. Type N shows a negative correlation, and types V and H show flattened distributions of data points with zero correlations. The vertically flattened distribution provides two leastsquare lines with the slopes ρ of 0 (blue) and −1 (orange), and the horizontally flattened distribution provides those with the slopes ρ of 0 (orange) and 1 (blue) in the REFER plots. Types P and P’ are positive correlations between log k and log k’. Type nr has no relation between log k and log k’, and the REFER plot may show a weak, artificial correlation trend.

**Fig. 3.**
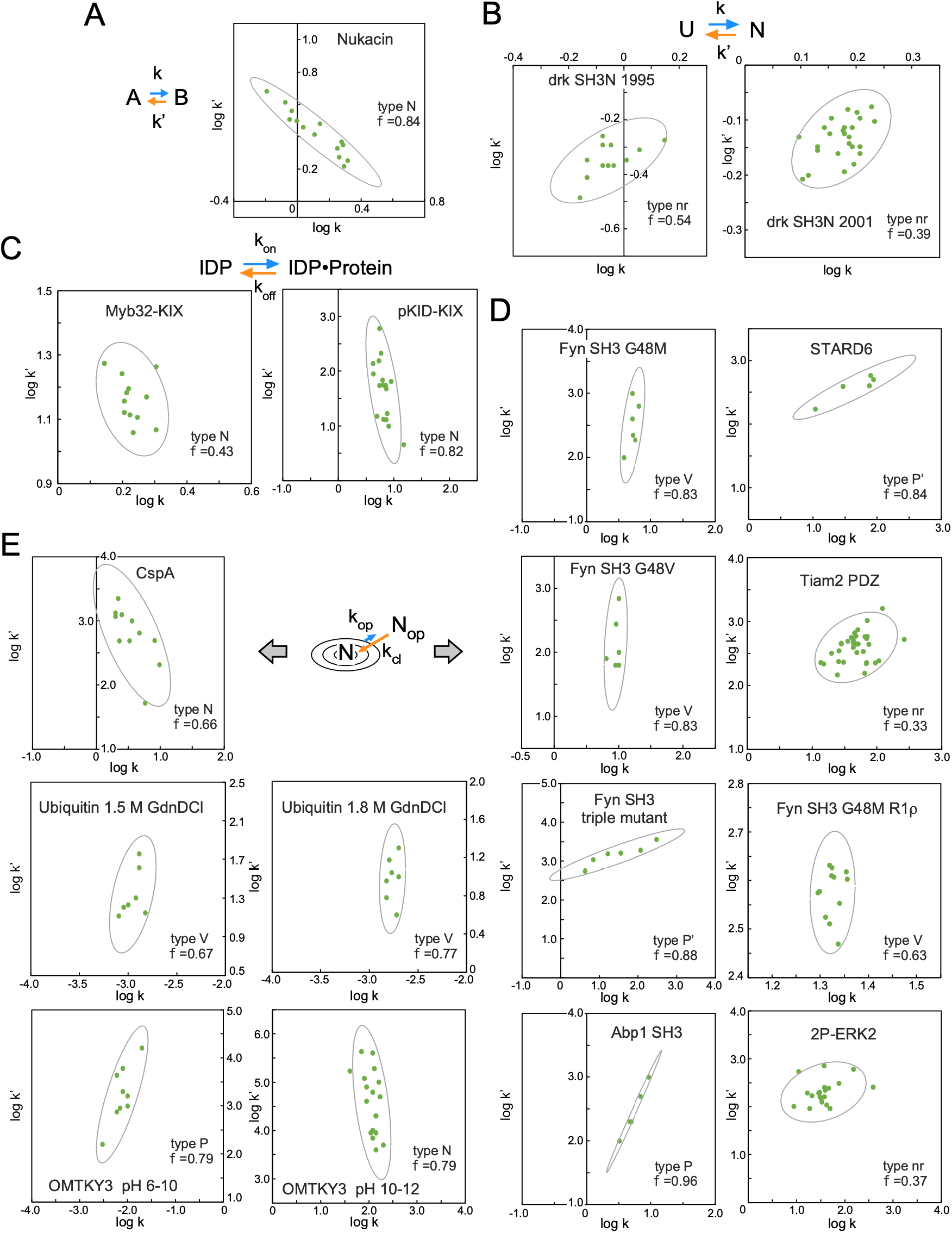
Datapoint distributions in the log k vs. log k’ plots. In each panel, protein/domain name, category type defined in Fig. 2, a 95% confidence ellipse, and the flatness of distribution are shown. The flatness of distribution is defined by f = 1-b/a, where a is the long axis and b is the short axis of the confidence ellipse. The axis ranges are set equally for the proper interpretation of the flatness of the confidence ellipses.

### Risk of overestimation of linearity

The imbalance in the numbers of experimental values to be obtained and parameters to be determined is a serious problem in the accurate and precise determinations of the residue-specific equilibrium and residue-specific kinetic rate constants. As for ^1^H-^15^N NMR, resonance- and state-specific NMR parameters, such as the relaxation rates, R_1_ and R_2_, of ^1^H and ^15^N nuclei, must be considered in addition to the objective equilibrium and kinetic rate constants. Global fitting is a solution, but only the average value over many residues is available. Alternatively, an appropriate assumption can be introduced to reduce the number of fitting parameters. For example, some parameters are supposed to remain unchanged under different measurement conditions (Table 1). Therefore, we have to pay attention to the risk that unexpected adverse effects caused by obligatory assumptions could generate an artificial linear relationship in REFER plots.

As already mentioned, the two axes of the REFER plot are not independent. Let ρ be the slope and a be the offset of rbLFER,

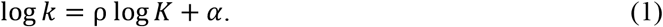

Let ε be the deviation factor of k due to the biases and errors,

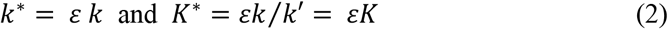

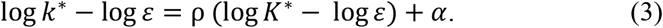

Equation 3 indicates that the data point moves by (log ε, log ε). This means that, if ρ has a value close to 0.5, the data point remains on the line and poses a risk of linearity overestimation.

In summary, the influences of measurement-specific biases and measurement errors must be considered seriously. The moderate linearity in the REFER plots does not simply prove a direct connection between the equilibrium constants and kinetic constants. The examination of the log k vs. log k’ plot may help to identify such misinterpretations. Ideally, a special insight from the REFER plot can be tested by the results obtained from other experiments. The retrospective analysis of apomyoglobin folding intermediates in the next section provides a clear illustration of this point.

### HX experiment of apomyoglobin

Information on the structural fluctuations of proteins can be obtained by monitoring the hydrogen/deuterium exchange of backbone amide protons with bulk water. The exchange mechanism consists of two processes: a two-state exchange of structural conversion and the exchange of the isotopes[32].

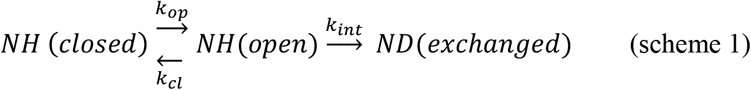

where NH(closed) is a folded state in which amide protons are protected from an exchange, and NH(open) is an open state in which exchange occurs. The H/D exchange rate, k_int_, is highly dependent on the pH of the solution. Usually, a single high pH pulse is used in the EX1 regime (k_int_ ≫ k_cl_) to simplify the analysis. Alternatively, a wide pH range of the labeling pulse can be used. Since the pH dependence of k_int_ is well known, k_op_ and k_cl_ can be determined without assuming the exchange mechanism, either EX1 (k_int_ ≫ k_cl_) or EX2 (k_int_ ≪ k_cl_). After a hydrogen/deuterium exchange reaction, NMR is used to measure the proton occupancy of each residue in an acidic solution. Mass spectrometry is also used after the fragmentation of proteins by protease digestion.

Sperm whale myoglobin is a popular model protein for understanding protein folding[33,34]. Myoglobin is a globular protein consisting of eight a-helices, designated A to H. The apo form of the protein has almost the same structure as the heme-bound holo form. Within the initial burst phase of apomyoglobin refolding, two kinetic intermediates, designated as I_a_ and I_b_, are sequentially formed[35]. In the state I_a_ structure, the major portions of helices A, G, and H and part of helix B are established. Subsequently, parts of helices C, D, and E are formed and added to the already-existing helices in the state I_b_ structure (see Fig. 5C)[33,34]. Wright’s group performed quenched-flow hydrogen exchange experiments using a continuous-flow mixer to determine the residue-specific kinetic parameters, k_op_ and k_cl_, of the folding intermediates[15]. The k_op_ and k_cl_ values were each assumed to remain unchanged in the two labeling pulse durations of 0.4-4.0 ms and 6.0-9.6 ms, and simultaneous numerical fitting was performed to obtain a more accurate estimation of the rate constants. Consequently, k_op_ and k_cl_ are averaged values of I_a_ and I_b_, respectively. We collected the k_op_ and k_cl_ data and associated errors from the literature[15] and constructed the REFER plot. We found that the (K, k_op_) and (K, k_cl_) data points were modestly aligned around straight lines (Fig. 4A). Upon closer inspection, several outlier data points were visually identified. Outlier residues (green and magenta) were removed to redraw the least-square lines, and the residues on the updated least-square lines formed a cluster of type N in the log k vs. log k’ plot (Fig. 4B and 4C). The slope ρ of the blue least-square line is 0.23. This is in contrast to the HX studies of the native states of other proteins, which provided almost zero slopes (Fig. 1D and 1E). This reflects the differences in the shapes of the energy surfaces. The intermediate states of protein folding are in a shallow basin, but the native state is located at the bottom of a deep basin (see Fig. 5B).

**Fig. 4.**
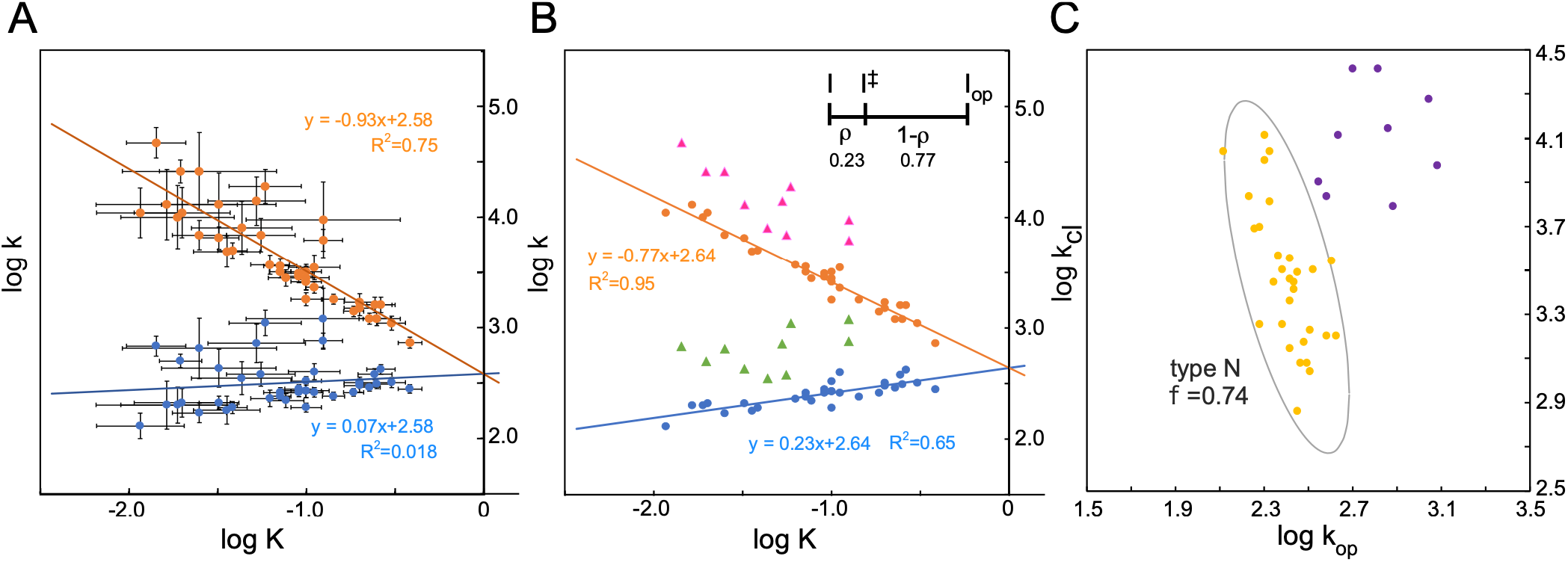
REFER plot and log k vs. log k’ plot of apomyoglobin folding intermediate. (A) REFER plot using all observed residues with estimated fitting uncertainties[15]. The data points and least square lines of the log k_op_ vs. log K plot and the log k_cl_ vs. log K plot are blue and orange, respectively. (B) Replot. Outlier residues (green and magenta) were removed to redraw the least-square lines (blue and orange). See Fig. S1 for details. The inset shows the position of the transition state I^‡^ on the reaction coordinate of the HX reaction. (C) Log k vs. log k’ plot. The yellow dots are the residues contributing to the least-square lines, and the purple dots are the outlier residues in (B). The yellow group belongs to type N. The yellow dots are enclosed by a 95% confidence ellipse.

**Fig. 5.**
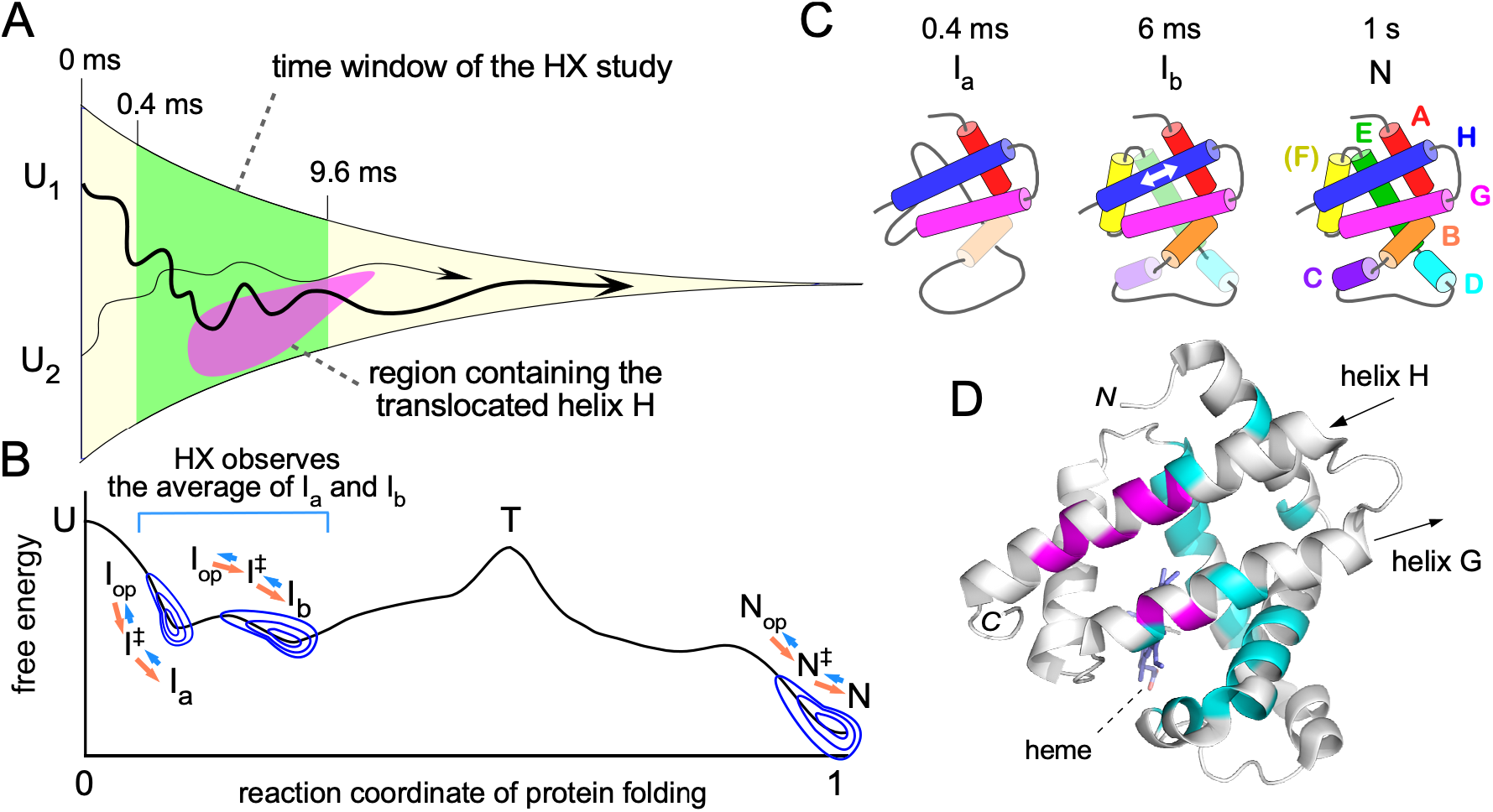
Non-native structure in the apomyoglobin folding intermediate. (A) Schematic of the folding funnel from the unfolded state U to the native state N. The thick and thin curved lines in the funnel represent individual folding paths from two initial states. The time window (0.4-9.6 ms) probed by the HX method is colored green. The conformational space containing the translocated helix H is colored magenta. The movement of the trajectory lines in and out of the magenta region illustrates the helix H translocation. (B) Energy diagram of the folding process. The concentric circles represent a basin-shaped energy landscape of the structural fluctuations in a microscopic thermodynamic equilibrium with the corresponding open state through the specific transition states, I^‡^ and N^‡^. T denotes the transition states of the entire folding process. (C) Sequential formation of the eight helices in apomyoglobin[33,34]. The kinetic intermediates I_a_ and I_b_ were observed at the 0.4 ms and 6 ms time points, respectively. The half-translucent cylinders represent partially formed a-helices. Helix F is not stably packed in the apo N state. The white arrow indicates the translocation of helix H. (D) Mapping of the outlier residues in the REFER plot on the N-state structure of myoglobin (PDB 2JHO). The native holo-structure is used as the best alternative to the intermediate structures. The residues on the least-square lines in Fig. 4B are colored cyan and the outlier residues are magenta. The other residues without information are colored white. These unprobed residues include parts of helices A, G, and H due to the full protection of the amide protons, and helix F and the loops due to the lack of local secondary structures in the intermediate states.

Figure 5 schematically shows the folding funnel and the energy diagram of apomyoglobin. The curved lines in the funnel depict refolding paths starting from unfolded states. The horizontal axis is the reaction coordinate of protein folding from 0 to 1. This axis can be regarded as the time axis, and an apomyoglobin molecule moves from left to right along a path. In the early time window (0.4 – 9.6 ms) from the start of refolding (green area in Fig. 5A), the intermediate states I_a_ and I_b_ are sequentially formed, and each amide group is in an exchange between the closed and open states, I_close_⇄I^‡^⇄I_open_, along the folding path (Fig. 5B). Note that the transition state I^‡^ in the HX exchange reaction is different from the transition state T in the entire refolding process, U → N.

The outlier residues were mapped on the N-state structure of apomyoglobin. All outlier residues (L104, F106, A134, L135, E136, L137, R139, D141, I142, and A143) are located at the interface between helix G and helix H (Fig. 5D, magenta). It is quite interesting that the Wright group showed that the intermediate I_b_ was a mixed state of two conformations, one of which contained the translocation of helix H by one helical turn toward its N-terminus relative to helix G (Fig. 5C), by using the HX study of amino acid mutants[16] and the combination of the HX study with fluorescence quenching and FRET (Förster resonance energy transfer) measurements[17]. The translocated helix H is not present in the folded state N, and thus they referred to the translocated form of helix H as a non-native structure. They also proposed that state I_b_ is in an energetic frustration at the B-G and G-H helix interfaces to facilitate the dynamic exchange of the two conformations. Strikingly, the region highlighted by the outlier residues precisely coincides with the amino acid residues involved in the non-native helix translocation (Fig. 5D, magenta)

## Discussion

In a two-state exchange process, in principle, the equilibrium constant and one of the two kinetic rate constants are independent, and the other kinetic rate constant is uniquely determined. Here, we showed that proteins have the property of the residue-based LFER (Fig. 1). The linear double logarithmic relationships between residue-specific equilibrium constants and the two kinetic rates of the same residue indicate that the thermodynamic and kinetic energy terms are closely related throughout a polypeptide chain. As a building block of a polymer, each amino acid residue of a protein participates in the conformational changes in a coordinated fashion to achieve smooth structural transformations. We consider this special feature to reflect a property of proteins acquired through evolution.

The residue-based LFER is an experimental demonstration of the consistency principle. This declaration can be proved by contradiction. Here, we use the term ‘non-native’ in the sense that it is absent in the two ground states including the unstructured state. If non-native structures or interactions are present in the transition state, then the involved amino acid residues can affect the free energy of the transition state independently of the ground states and the position of the corresponding data point will deviate from the least square line. In rare instances, however, the data points are aligned by chance. Thus, in a precise mathematical sense, the role of rbLFER is more of a statistical null hypothesis test than a rigorous mathematical proof of the consistency principle. In other words, it is almost impossible to explain the physicochemical basis of the linear relationships in Fig. 1 by the assumption that the transition states contain non-native structures or interactions.

The consistency principle represents an ideal protein. Real proteins do possess some non-ideality due to minimal frustrations. An ideal protein would provide (K, k) data points on a perfectly straight line in the REFER plot, whereas a real protein provides (K, k) data points scattered around a straight line, even after perfect measurement error removal. Here, we found the deviations from the least-square lines in the REFER plot for the apomyoblobin folding intermediates (Fig. 4). The distribution of the outlier residues in the three-dimensional structure is in good agreement with the non-native translocation of helix H in intermediate I_b_, which was previously identified by other methods (Fig. 5D). The prevention of the formation of t he translocated form by an engineered disulfide bond increased the overall folding rate by two-fold[17], and the stabilization of the translocated form by mutations in helix F decreased the folding rate down to 20%[34]. These results are in agreement with the consistency principle: a non-native structure on the folding pathway delays the overall folding process.

The quenched-flow HX experiment of apomyoglobin examined the structural conversion between I_close_ and I_open_ through the transition state I^‡^ along the folding path (Fig. 5B). For better precision, the two exchange processes, I_a_ ⇄ I^‡^ ⇄ I_op_ and I_b_ ⇄ I^‡^ ⇄ I_op_, were jointly analyzed to provide their averaged picture[15]. First, we focus on the structural fluctuation of I_b_. The I_op_ state connected with I_b_ is more or less similar to I_a_. We assume that I^‡^ contains a greater fraction of the translocated helix H than that in the ground state I_b_. The increase in the fraction of the translocation is the ‘non-native structure’ that is absent in the two ground states, I_b_ and I_op_. Another interpretation is the elevated exchange rate of the helix translocation. In this case, the increase in the rate is the ‘non-native interaction’ that is absent in the two ground states. These non-native structures/interactions violate the consistency principle and generate the outlier residues in the REFER plot. In contrast to I_b_, I_a_ contains fewer secondary structures and thus does not contribute much to the deviation of the REFER plot. In sum, the rbLFER of the HX study detected non-native structures/interactions on the folding pathway by observing the transition state I^‡^ (Fig. 5B).

## Conclusion

The linear relationship between log K (the logarithm of the equilibrium constant) and log k (the logarithm of the rate constant) in the REFER plot holds for structural changes of many proteins, according to our retrospective analysis (Fig. 1). The analytical methods include EXSY NMR, R_2_ dispersion relaxation NMR, and HX measurements. The residue-based LFER is seen in the two-state slow exchange (Fig. 1A) and structural fluctuations (Fig. 1D and 1E) of monomeric small proteins, and in two-molecule systems, such as IDP-protein interactions (Fig. 1C). The substantial number of rbLFER examples indicates the applicability of the consistency principle to a wide variety of protein-related phenomena. The linearity of the REFER plot can be used as a measure of the deviation from ideal conditions for smooth protein structural changes. The reanalysis of the HX study of apomyoglobin[15] is particularly fruitful (Figs. 4 and 5). This unexpected outcome demonstrates that the rbLFER is not a special experiment to prove the consistency principle but is a practical method to obtain dynamic information of proteins. Finally, one must bear in mind that measurement biases and errors could lead to the overestimation of linearity in the REFER plots. The development of a novel methodology is keenly anticipated for bias-free, accurate determinations of the residue-specific exchange parameters of polypeptide chains.

## Methods

We performed a literature search to collect residue-specific equilibrium and residue-specific kinetic constants of proteins, mainly in the PubMed literature database (http://www.ncbi.nlm.nih.gov/pubmed/). The keywords included ‘two states’, ‘two sets of cross peaks’, ‘exchange spectroscopy’, ‘residue-specific’, LFER, etc., and their combinations. The linear regression analyses of the REFER plots and log k vs. log k’ plots were performed in the Excel files. The protein cartoon was generated with the program PyMOL, version 2.4.2 (Schrödinger). The cartoon image of the apomyoglobin was generated using the PDB entry 2JHO.

## Abbreviations used

CPMG: Carr-Purcell-Meiboom-Gill
EXSY: exchange spectroscopy
FRET: Förster resonance energy transfer
GdnDCl: Guanidine deuterium chloride
HSQC: heteronuclear single quantum coherence
HX: hydrogen exchange
IDP: intrinsically disordered polypeptide
LFER: linear free energy relationship
rbLFER: residue-based LFER
RD: relaxation dispersion
REFER: rate-equilibrium free energy relationship

## Acknowledgments

This work was supported by the Japan Society for the Promotion of Science (JSPS, Japan), KAKENHI Grant Number JP21H02448, and the Mitsubishi Foundation (Japan) Research Grants in the Natural Sciences, Grant Number 202110017, to D.K.

## Author Contributions

Conceptualization, D.F., and D.K.; Investigation, D.F., H.S., and D.K.; Writing – Original Draft, D.K.; Writing – Review & Editing, F.D., and S.H.; Funding Acquisition, D.K.

## Appendix A. Supplementary data

Supplementary data for this article include one figure (PDF).

Details of the REFER plot of the apomyoglobin folding intermediate (PDF)

---

**Fig. S1.**
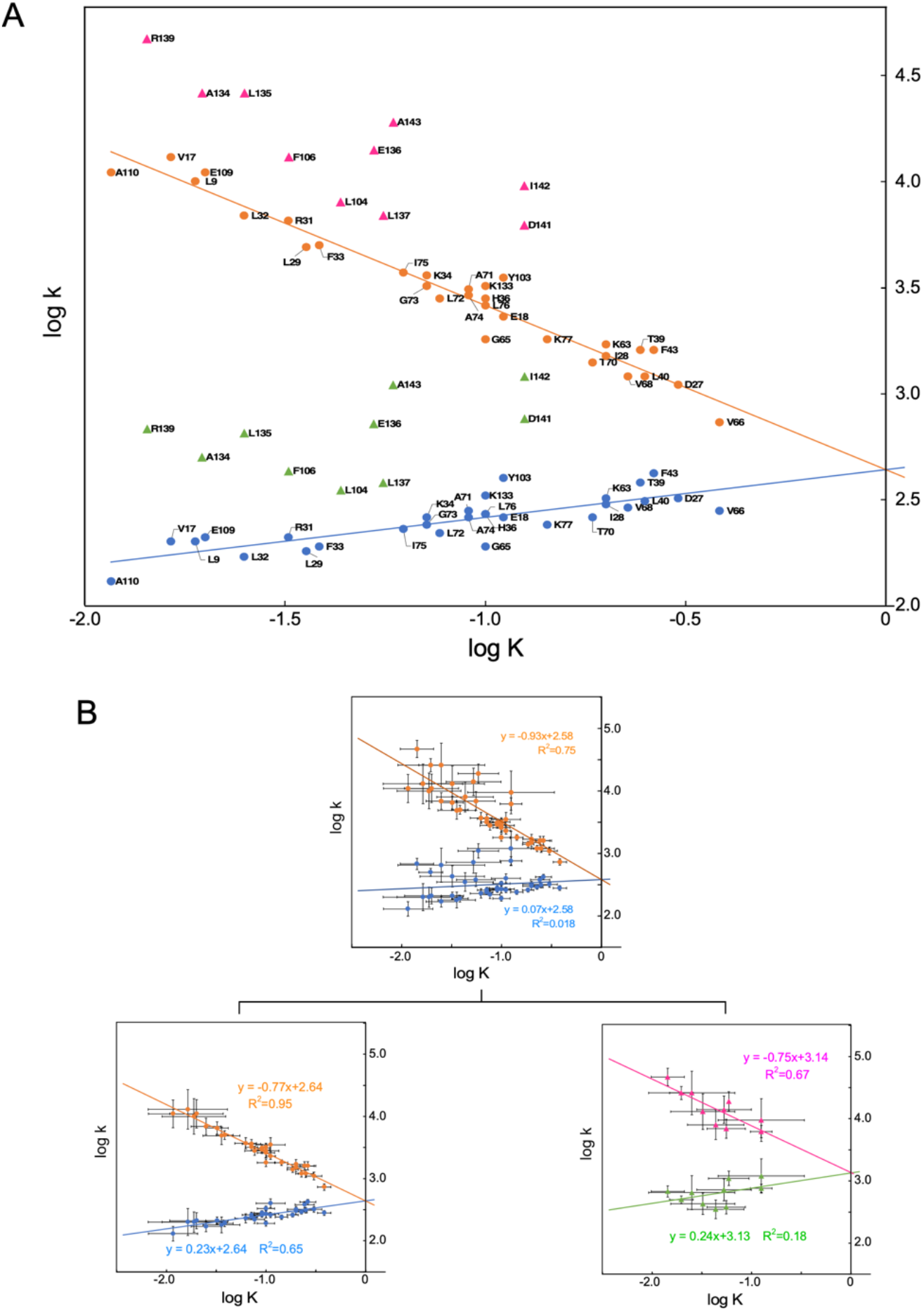
Details of the REFER plot of the apomyoglobin folding intermediate. (A) Labeling of amino acid residues. (B) Error bars of the first least-square lines (blue and orange) and the second least-square lines (green and magenta). The second lines are almost parallel to the first lines, which might suggest a hidden mechanism for the formation of the non-native structure, but the large errors hamper proper interpretation.

